# Memory precision of object-location binding is unimpaired in *APOE* ε4-carriers with spatial navigation deficits

**DOI:** 10.1101/2020.12.18.423245

**Authors:** Helena M. Gellersen, Gillian Coughlan, Michael Hornberger, Jon S. Simons

**Author notes:** Corresponding author Department of Psychology, University of Cambridge, Downing Street, Cambridge CB2 3EB.

## Abstract

Research suggests that tests of memory fidelity, feature binding and spatial navigation are promising for early detection of subtle behavioural changes related to Alzheimer’s disease (AD). In the absence of longitudinal data, one way of testing the early detection potential of cognitive tasks is through the comparison of individuals at different genetic risk for AD. Most studies have done so using samples aged 70 years or older. Here, we tested whether memory fidelity of long-term object-location binding may be a sensitive marker even among cognitively healthy individuals in their mid-60s by comparing participants at low and higher risk based on presence of the ε4-allele of the apolipoprotein gene (*n*=26 ε3ε3, *n*=20 ε3ε4 carriers). We used a continuous report paradigm in a visual memory task that required participants to recreate the spatial position of objects in a scene. We employed mixture modelling to estimate the two distinct memory processes that underpin the trial-by-trial variation in localisation errors: retrieval success which indexes the proportion of trials where participants recalled any information about an object’s position and the precision with which participants retrieved this information. Prior work has shown that these memory paradigms that separate retrieval success from precision are capable of detecting subtle differences in mnemonic fidelity even when retrieval success could not. Nonetheless, a Bayesian analysis found good evidence that ε3ε4 carriers did not remember fewer object locations (*F*(1, 42)=.450, *p*=.506, BF_01_ =3.02), nor was their precision for the spatial position of objects reduced compared to ε3ε3 carriers (*F*(1, 42)=.12, *p*=.726, BF_01_ =3.19). Because the participants in the sample presented here were a subset of a study on *APOE* effects on spatial navigation in the Sea Hero Quest game (Coughlan et al., 2019. *PNAS, 116*(9)), we obtained these data to contrast *APOE* effects on the two tasks within the same sample (*n*=33). Despite the smaller sample size, wayfinding deficits among ε3ε4 could be replicated (*F*_(1, 33)_ =5.60, *p*=.024, *BF*_*10*_ =3.44). Object-location memory metrics and spatial navigation scores were not correlated (all *r*<.25, *p*>.1, 0<BF_10_ <3). These findings show spared object-location binding in the presence of a detrimental *APOE* ε4 effect on spatial navigation. This suggests that the sensitivity of memory fidelity and binding tasks may not extend to individuals with one ε4-allele in their early to mid-60s. The results provide further support to prior proposals that spatial navigation may be a sensitive marker for the earliest AD-dependent cognitive changes, even before episodic memory.

## 1. Introduction

Alzheimer’s disease (AD) has a long preclinical phase during which pathological neural changes occur without overt, detrimental effects on behaviour (Jack *et al*., 2010; Sperling *et al*., 2011; Jack and Holtzman, 2013; Sutphen *et al*., 2015). This long preclinical phase offers the possibility of interventions that may target further pathological changes and prevent irreversible cell death (Chetelat *et al*., 2010; Rentz *et al*., 2013). Cognitive tests are the most cost-effective and simple way to screen for cognitive impairment related to dementia, yet standard neuropsychological tests typically fail to detect these subtle preclinical symptoms of AD pathology (Salmon, 2011; O’Donoghue *et al*., 2018). In the absence of longitudinal data, individuals with high risk for late-onset AD based on the ε4-allele of the apolipoprotein (*APOE*) gene are a good model to test the diagnostic sensitivity of cognitive tests because they are more likely than ε3ε3 carriers to develop the disease, exhibit AD pathology at an earlier point in time and decline at a more rapid rate (Corder *et al*., 1993; Raber *et al*., 2004; Caselli *et al*., 2011; Caselli and Reiman, 2013; Risacher *et al*., 2013; Grilli *et al*., 2018; Flowers and Rebeck, 2020). *APOE* ε4-carriers exhibit deficits in tests of long-term feature binding, mnemonic fidelity and spatial navigation, making these tasks promising markers of incipient cognitive decline related to AD (Coughlan *et al*., 2018; Stark *et al*., 2019; Zokaei *et al*., 2019). Yet, there are few studies testing these tasks in neuropsychologically unimpaired middle-aged ε4-carriers, and even fewer studies looked at more than one of these different types of tasks in the same sample. Here, we determined whether a novel test of long-term object-location binding is sensitive to *APOE* effects in a sample that previously exhibited spatial navigation deficits (Coughlan *et al*., 2019).

Older adults, individuals with mild cognitive impairment (MCI) and preclinical individuals with positive AD biomarkers exhibit significant deficits in mnemonic discrimination of novel and studied targets under conditions of high feature overlap (Trelle *et al*., n.d.; Yassa *et al*., 2010, 2011; Ally *et al*., 2013; Stark *et al*., 2013; Reagh *et al*., 2014; Stark and Stark, 2017; Berron *et al*., 2018, 2019; Leal and Yassa, 2018; Gellersen *et al*., 2020; Webb *et al*., 2020). Similarly, cognitively healthy preclinical adults (defined by *APOE* genotype or AD pathologies), as well as MCI and AD patients, also perform significantly worse in tests of feature binding, showing a marked decline in representational fidelity (Atienza *et al*., 2011; Rentz *et al*., 2011; Troyer *et al*., 2012; Della Sala *et al*., 2012; Hampel, 2013; Bastin *et al*., 2014; Oedekoven *et al*., 2015; Parra *et al*., 2015, 2019; Van Geldorp *et al*., 2015; Koppara *et al*., 2015; Mowrey *et al*., 2016; Pietto *et al*., 2016; Chen and Chang, 2016; Liang *et al*., 2016; Polcher *et al*., 2017; Zokaei *et al*., 2019; Delhaye *et al*., 2019; Konijnenberg *et al*., 2019; Pavisic *et al*., 2020; Valdés *et al*., 2020; Korkki *et al*., 2020).

We capitalise on current evidence for subtle cognitive deficits in preclinical AD by using a memory precision task with demands on memory binding and fidelity of mnemonic representations, abilities that are particularly affected by AD pathology even from preclinical stages onwards (Rentz *et al*., 2013; Ritchie *et al*., 2017; Berron *et al*., 2019). We use study-test delays that preclude the use of short-term memory. *APOE* ε4-carriers have an advantage in short study-test delays but may be predisposed to accelerated rate of forgetting thereafter (Zokaei *et al*., 2019; Pavisic *et al*., 2020). A longer study-test delay may be able to index such faster forgetting. We hypothesised that our task design may detect ε4-dependent differences because 1) the task involves entorhinal and hippocampal mediated relational binding of objects and locations, which is impaired in prodromal AD (Charles *et al*., 2004; Reagh *et al*., 2014; Hampstead *et al*., 2018; McIlvain *et al*., 2018; Weigard *et al*., 2020), 2) a continuous metric may be a more sensitive index than categorical measures of retrieval (Zokaei *et al*., 2015; Korkki *et al*., 2020), and 3) memory fidelity relies on communication between hippocampus and cortical regions, which exhibit altered connectivity in the early course of AD (Buckner *et al*., 2005; Sperling *et al*., 2011; Jack *et al*., 2015; Richter *et al*., 2016; Xie, 2018; Stevenson *et al*., 2018; Cooper and Ritchey, 2019; Harrison *et al*., 2019; Sullivan *et al*., 2019; Berron *et al*., 2020; Foo *et al*., 2020).

Only one study has tested the fidelity of relational binding with longer memory retention intervals using continuous report paradigms in an *APOE* genotyped cohort in their 60s (Zokaei *et al*., 2019), showing a reduction in the fidelity of object-location binding for preclinical ε4 homozygote older adults. No such effect for was present in ε3ε4 heterozygotes when using the mean error between target location and response as a performance metric. The presence of an effect of the ε4-allele on the fidelity of long-term object-memory binding is promising as it suggests that this task is potentially sensitive to preclinical AD-related changes even in individuals in their 60s. Performance reductions might be observed not just in ε4 homozygotes but also heterozygotes for object-location binding when using a more sensitive index than mean localisation error, such as localisation precision which controls for accessibility of any information from memory. Another option may be to increase interference by adding more objects to studied scenes to place further demands on transentorhinal and hippocampal processes, (Kirwan and Stark, 2007; Newsome *et al*., 2012; Reagh and Yassa, 2014) thereby resulting in more misbinding errors among individuals with poorer mnemonic representations (Liang *et al*., 2016; Hampstead *et al*., 2018). Here, we use both approaches to investigate whether continuous report paradigms can be made even more sensitive to AD risk.

We examine the utility of this novel test of memory fidelity of relational binding that engages regions vulnerable to early AD, supplemented with a mixture modelling approach that specifically indexes precision, to test the effect of the ε4-allele on the precision of object-location binding beyond short-term memory retention. We compare model-derived metrics with those used in prior studies with continuous report paradigms such as those by Zokaei et al. (2019) to determine if the separation of precision and retrieval success may be able to tease apart subtle *APOE* effects on memory abilities. We apply our test to a sample that has previously been characterised in terms of spatial navigation abilities (Coughlan *et al*., 2019). An added benefit of our study is therefore to test whether a fidelity metric for spatial memory will be similarly sensitive to AD risk as spatial navigation measures. To our knowledge, no other study to date provides data on the effect of the ε4-allele on spatial memory fidelity and spatial navigation in the same *APOE*-genotyped sample.

## 2. Methods and methods

### 2.1 Participants

The study was carried out at the University of East Anglia, Norwich with ethical approval from the Faculty of Medicine and Health Sciences Ethics Committee at UEA (Reference FMH/2016/2017-11). Exclusion criteria were cognitive impairment and neuropychiatric conditions. Participants provided written informed consent before participation. The sample presented here was previously described by Coughlan and colleagues (2019). The sample size was based on that of prior studies that investigated the effect of the ε4-allele on spatial navigation (Kunz *et al*., 2015).

Forty-nine participants completed the spatial precision memory task. We included *n*=26 individuals with the ε3ε3 genotype aged 53 to 74 (*M*=63.38, *SD*=6.07; 13 females) and *n*=20 individuals with the ε3ε4 genotype aged 54 to 80 (*M*=64.80, *SD*=6.83; 5 females) for our main analysis. Three volunteers with the ε4ε4 genotype aged 63 or 64 years also completed the test battery (*M*=63.33, *SD*=.58; 1 female). Given the small number of ε4 homozygotes and the differences between ε3ε4 and ε4ε4 carriers in general, our main analysis focused on a comparison of ε3ε3 carriers and ε4 heterozygotes to avoid the admixture of different genotypes. In a sensitivity analysis, we determined whether the addition of the high risk ε4 homozygotes influenced the results. Sample demographics and standard neuropsychological scores are shown in Table 1.

**Table 1.** Demographics and standard neuropsychological test scores by *APOE* genotype group.

|  | ε3ε3<br>(*n*=26)<br>*Mean (SD)* | ε3ε4<br>(*n*=20)<br>*Mean (SD)* | ε4ε4<br>(*n*=3)<br>*Mean (SD)* |
| --- | --- | --- | --- |
| **Age** | 63.4 (6.07) | 64.8 (6.83) | 63.3 (.58) |
| **Sex** |  |  |  |
| Female | 13 (50%) | 5 (25%) | 1 (33%) |
| Male | 13 (50%) | 15 (75%) | 2 (67%) |
| **Education** | 14.25 (2.31) | 13.80 (2.26) | 14.67 (.58) |
| **ACE Total** | 93.9 (4.91) | 94.6 (2.42) | 93.0 (3.61) |
| **ACE Memory** | 25.0 (1.50) | 25.1 (1.00) | 24.7 (1.53) |
| **ROCF immediate** | 33.3 (2.59) | 31.8 (2.69) | 32.0 (2.65) |
| **ROCF delayed** | 21.4 (5.73) | 21.9 (4.67) | 22.2 (9.78) |

Note: ACE: Addenbrookes Cognitive Examination; ROCF: Rey-Osterrieth Complex Figure. Delayed copy was three minutes after presentation. The genotype groups did not differ significantly in terms of age ( $F_{(2,46)}=.30$ , $p=.739$ ) or scores on the Addenbrookes Cognitive Examination (ACE) regardless of whether the total score ( $F_{(2,46)}=.11$ , $p=.90$ ) or the memory sub-score was used ( $F_{(2,46)}=.28$ , $p=.760$ ).

In this sample, Coughlan and colleagues previously tested spatial navigation performance at two time points (Coughlan *et al*., 2019, 2020). Sixty participants (29 ε3ε3, 31 ε3ε4) completed the SHQ at baseline. At follow-up, 59 remained to complete the spatial navigation tasks, 49 of whom were also given the precision memory task and are included in this study. We then compared the spatial navigation data from the baseline assessment with our object-location precision memory task from the follow-up session. Although this has the caveat that the spatial navigation data were obtained 18 months prior to the memory data, we decided that the issue of practise effects at re-test was a greater confound because it could have allowed participants to develop strategies to better cope with the demands the spatial navigation task. In their test-retest analysis, Coughlan et al. (2020) suggest that this may have indeed been the case and that the reduction of novelty in the spatial navigation task may reduce its diagnostic utility because poor performers improved more than those with initially better scores. Using the first assessment of both memory and spatial navigation tasks is therefore more informative to determine whether effects of *APOE* can be observed in each cognitive function.

### 2.2 Precision memory task

Details of the precision memory task can be found in the Supplementary Material. Briefly, participants were asked to remember the identity and locations of objects in a scene. Each encoding display consisted of a trial-unique background image with three objects pseudorandomly arranged around an invisible circle centred at the midpoint of the image. Object positions were constrained to maintain a minimum of 62.04° between objects to avoid spatial overlap. Participants undertook five practice trials before beginning the actual task. The main task comprised five study-test blocks. In each of the five blocks, participants first viewed five displays during the study phase. After encoding, an interference task required participants to count backwards from a random number between 50 and 100 for 12 seconds to prevent active rehearsal of memorised displays. Each test trial began with an identification question where participants were asked to determine which of two presented objects had previously been shown. If they chose correctly, the associated background image appeared, and participants were asked to move the object around the screen to recreate its studied location as precisely as possible. Participants viewed 25 displays and completed 75 test trials, each containing an identification and a localisation question.

### 2.3 Spatial navigation task

To compare the effects of *APOE* on the object-location memory task and spatial navigation in this same sample, we obtained the previously published spatial navigation data (Coughlan *et al*., 2019, 2020), from the Sea Hero Quest (SHQ) app (Coutrot *et al*., 2018).The SHQ has previously been described in detail. Briefly, SHQ is a game in which participants navigate through a virtual environment to reach checkpoints described on a map they study at the beginning of each level. Crucially, the maps are shown in an allocentric perspective but once a level begins, participants navigate based on an egocentric viewpoint. Participants played three different levels. Performance metrics were wayfinding distance and average distance to the border of an environment to index border bias (Coughlan *et al*., 2019).

### 2.4 APOE genotyping

DNA was collected using a Darcon tip buccal swab (LE11 5RG; Fisher Scientific). Buccal swabs were refrigerated at 2–4 °C until DNA was extracted using the QIAGEN QIAamp DNA Mini Kit (M15 6SH; QIAGEN). DNA was quantified by analyzing 2-μL aliquots of each extraction on a QUBIT 3.0 fluorometer (LE11 5RG; Fisher Scientific). DNA extractions were confirmed by the presence of a DNA concentration of 1.5 μg or higher per 100 μg of AE buffer as indicated on the QUBIT reading. PCR amplification and plate read analysis was performed using Applied Biosystems 7500 Fast Real-Time PCR System (TN23 4FD; Thermo Fisher Scientific). TaqMan Genotyping Master Mix was mixed with two single-nucleotide polymorphisms of *APOE* (rs429358 at codon 112 and rs7412 at codon 158). These two single-nucleotide polymorphisms determine the genotype of *APOE*2, Ε3, and Ε4 (2007; Applied Biosystems).

### 2.5 Statistical analysis

#### 2.5.1 Mixture modelling

Models fitted to the data and distribution of responses across all participants by genotype are shown in Fig. 2. We fit probabilistic mixture models to the location placement errors expressed as the degrees separating the response from the target (Zhang and Luck, 2008; Bays *et al*., 2011; Suchow *et al*., 2013; Richter *et al*., 2016; Zokaei *et al*., 2020). The approach aims to determine the distribution of trial responses in order to examine which retrieval mechanisms best explain the observed responses: i) correctly recalled locations, ii) random guesses or iii) a misbinding error in which the location of the target is confused with that of another object from the same display. Guess trials were modelled using a uniform distribution. The proportion of trials within the uniform distribution represents the guess rate *pU* and 1-*pU* expresses retrieval success *pT*. Correctly remembered items were modelled by a circular Gaussian (von Mises) distribution centred at the target location with its standard deviation reflecting the precision with which locations are recalled. Larger standard deviations (*SD*) correspond to lower localisation fidelity. Misbinding errors were modelled by von Mises distributions centred around the two distractor items.

**Figure 1.**
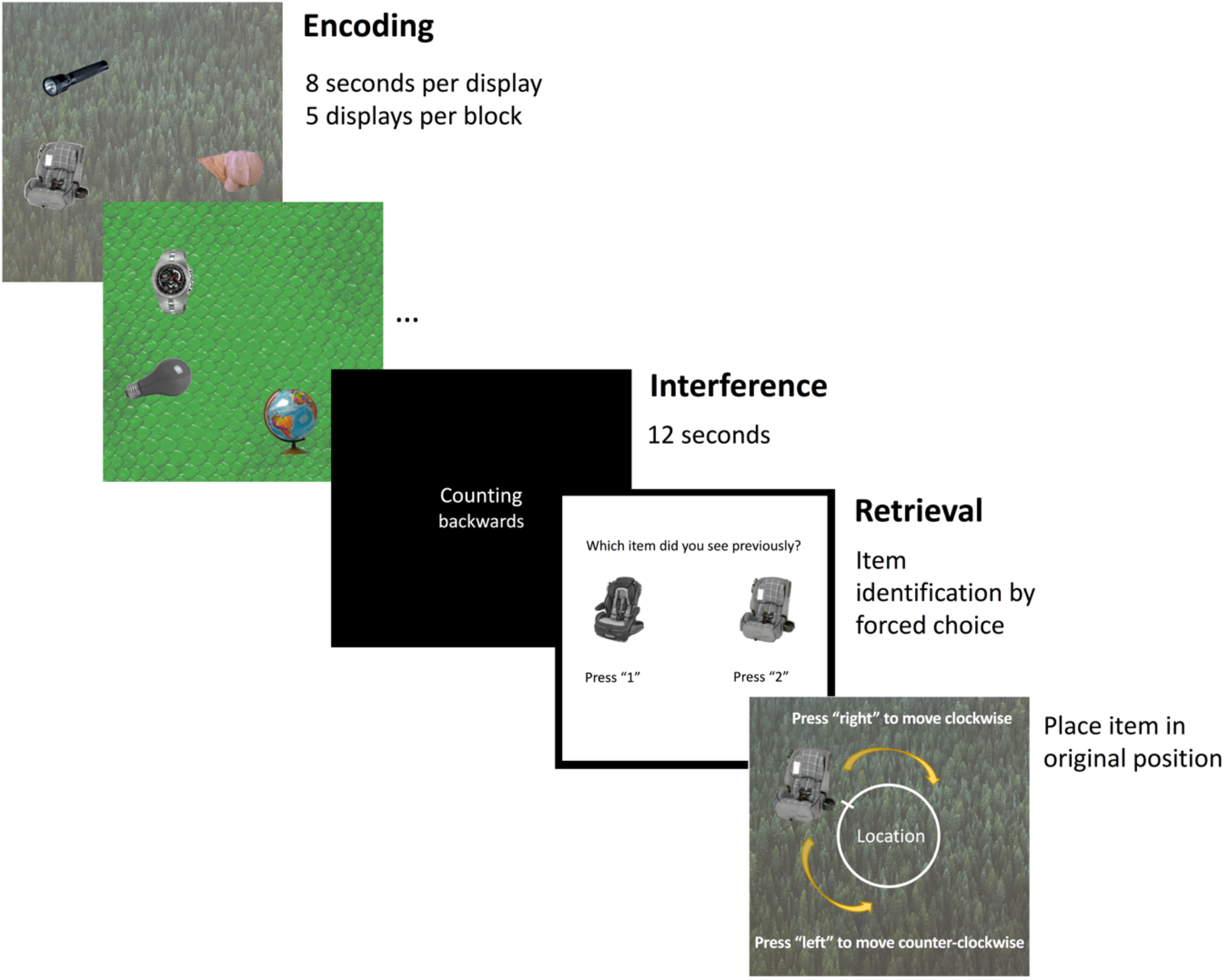
Schematic of the precision memory task.

**Figure 2.**
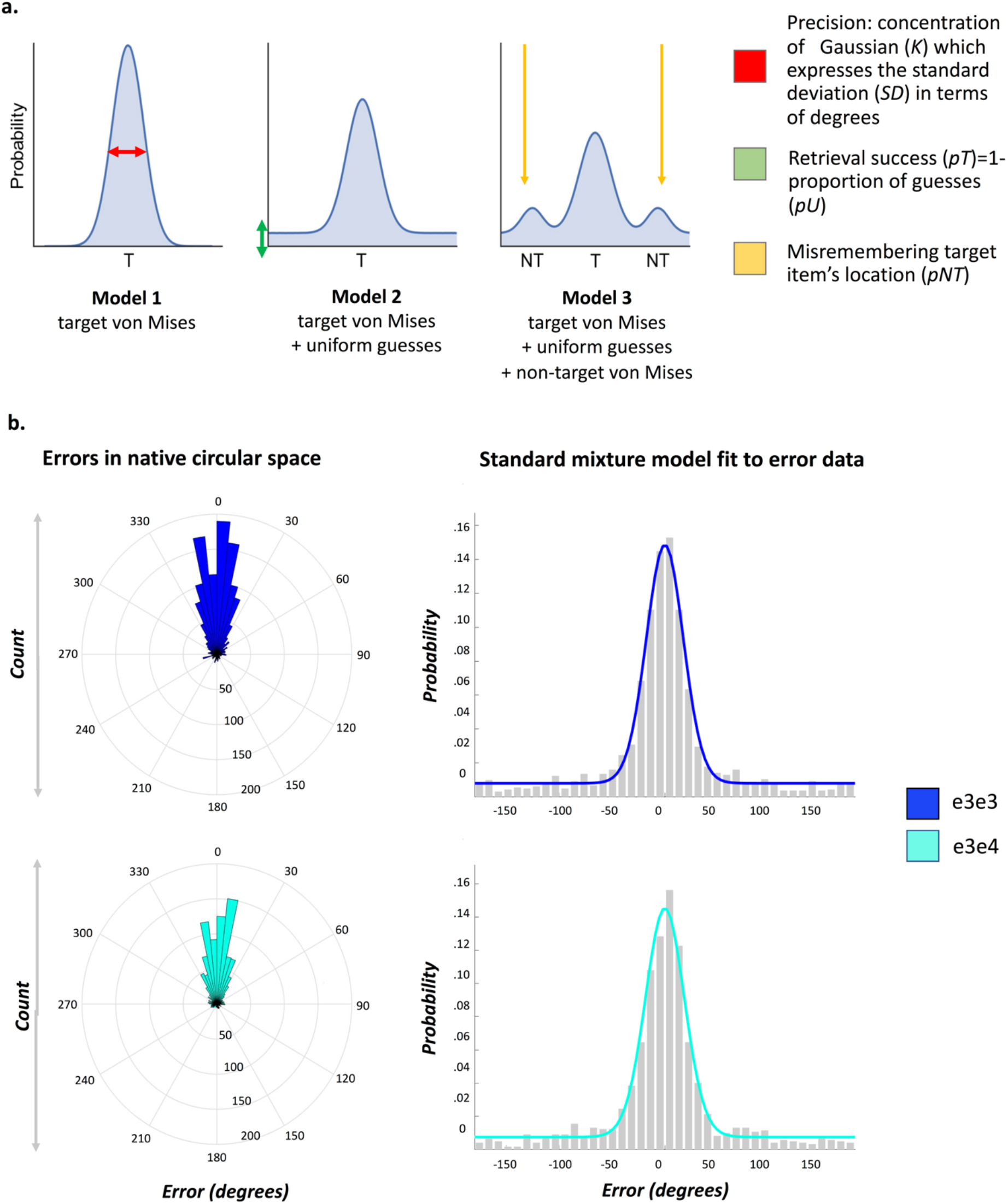
Tested models and results from the mixture modelling approach. (a) Proposed models to capture location memory performance. In Model 1, all object locations are assumed to be correctly recalled without any guess responses (probability of guessing: *pU*=0). The mean distance of responses from the target can be represented by the width of the von Mises (circular Gaussian) distribution, expressing the precision of memory recall (expressed as the standard deviation (*SD*) of the von Mises distribution where higher values reflect less precision; for a more intuitive interpretation where higher values reflect better performance, the *SD* value can be converted to the von Mises distribution concentration parameter *K*; see Supplementary Material). Model 2 assumes a mixture of guessed and correctly remembered responses, where the proportion of responses that fall within the uniform distribution is denoted by the parameter *pU* that captures the proportion of guessed responses. For a more intuitive understanding where higher values reflect higher performance this parameter can also be expressed as retrieval success denoted by *pT*, the proportion of trials within the von Mises distribution, i.e. trials in which the target location was correctly recalled. Model 3 assumes that responses reflect a combination of guessing, correctly remembered responses with variable degree of precision, and swaps of target and distractor locations, represented as von Mises distributions centered at the locations of distractor objects. **(b) Distribution of location errors by ε4-status in native circular space (left hand side) and the Standard Mixture Model (von Mises + uniform) fit to responses**. Model 2 was identified as the best fitting model in a model comparison procedure detailed in the Supplementary Material.

To maximise comparability with the only other study on the effect of the *APOE* genotype on location memory precision (Zokaei *et al*., 2020), we also used Bayesian modelling implemented with the MemToolbox in MATLAB 2016a (Suchow *et al*., 2013). We fit three models to the error data collapsed across all participants by *APOE* genotype group to test whether which components could explain localisation performance. The models contained the following components (Fig. 2A): Model 1 (von Mises distribution) assumes that no guessing occurred; Model 2 (uniform + von Mises distribution) assumes that responses reflect a mixture of guessed trials and correctly recalled locations with response-to-target distance varying across trials; Model 3 (uniform + von Mises + von Mises for non-targets) extends Model 2 by assuming that some incorrect responses were due to object-location misbinding. Deviance Information Criterion favoured Model 2. All further analyses are conducted using this model. For more details on modelling and comparison with an alternative model fitting procedure based on work by Bays and colleagues (Richter *et al*., 2016; Korkki *et al*., 2020) refer to the Supplementary Material.

This approach allowed us to test if the *APOE* ε4-allele affects the probability of correctly retrieving information from memory and/or mnemonic fidelity (i.e. precision with which this information is recalled). Mixture modelling is superior to other approaches that distinguish between retrieval success and fidelity of retrieved information because the estimation of the uniform distribution accounts for guess responses placed near the target item.

We calculated retrieval success and precision for each subject. To improve robustness of estimates for precision and retrieval success, we calculated a cut-off for guessing from the mixture modelling approach across the full sample following the examples of prior studies (for details see Supplementary Material) (Richter *et al*., 2016; Korkki *et al*., 2020). Localisation errors exceeding 63° response-to-target distance were deemed as failure to retrieve an object’s location. For each subject we calculated retrieval success as the proportion of trials with errors ≤63°. A measure of imprecision was derived from the standard deviation across all responses with localisation errors ≤63°.

#### 2.5.2 APOE group differences memory performance

We employed a combination of frequentist (two-tailed tests with a statistical significance level of *p*<.05) and Bayesian methods to test for *APOE* genotype effects.

##### 2.5.2.1 Mixture modelling by APOE genotype group

We first tested for differences in guessing (*g*) and imprecision (*SD*) estimates for the standard mixture models fit to all responses across subjects in the ε3ε3-carrier and ε3ε4-carrier group, respectively. To obtain a *p*-value, true group differences were compared to the distribution of standardised differences obtained from random group assignments over 1000 permutations (sample 1 with *n*=26 to match the number of participants in the ε3ε3 group; sample 2 with *n*=20, as in the ε3ε4 group). This approach has the advantage of operating on more robust model parameters due to reduced noise resulting from larger number of trials available for mixture modelling.

##### 2.5.2.2 APOE effects based on single-subject scores

Next, we carried out analyses on individual subject data while controlling for nuisance variables using a linear model with sex and age as covariates and *APOE* genotype as between-subjects factor of interest. Dependent variables were the proportion of correctly identified items, and the measures of retrieval success and precision. Cohen’s *f*^*2*^ was used to denote the effect size of the *R*^*2*^-change from a model with covariates (age, sex) to a model with *APOE* genotype (ε3ε3 vs. ε3ε4). We also calculated the Bayes Factor for the contrast of the model with covariates and the full model with covariates and *APOE* genotype as between-subjects factor of interest using the R package *BayesFactor* (https://CRAN.R-project.org/package=BayesFactor). A Bayes Factor of >3 was deemed as good evidence in support of the alternative hypothesis if indexed by *B*_*10*_ and for the null hypothesis if indexed by *B*_*01*_ (Jeffreys, 1961; Keysers *et al*., 2020; Korkki *et al*., 2020).

##### 2.5.2.3 Supplementary analyses for precision memory

In order to make our results more comparable with prior studies that used a similar object-location binding paradigm without mixture modelling to separate retrieval success from precision (Zokaei *et al*., 2017, 2019), we also calculated the mean absolute error between targets and responses to determine whether a modelling approach to separate retrieval success and memory precision may be more sensitive to detect *APOE* effects.

We conducted a control analysis termed ‘nearest neighbour analysis’ as used in prior work (Pertzov *et al*., 2013; Zokaei *et al*., 2019). This analysis allowed us to test whether there was a difference in the nature of incorrect responses between genetic groups by considering the occurrence of misbinding errors. A significant *APOE* effect on the distance to the nearest neighbour would suggest that error responses in the two groups are not caused by the same mechanisms. The group with significantly smaller nearest neighbour difference is likely to commit more misbinding errors.

Prior work has demonstrated an interaction between study-test delay and the *APOE* ε4-allele on short-term memory versions of continuous object-location tests with ε4-carriers at an advantage at short delays of 1s which subsides at longer delays beyond 4s (Zokaei *et al*., 2017, 2020). Using the correlation between localisation error and study-test delay in each subject, we tested whether ε4-carriers exhibit steeper performance decline as a function of delay.

Finally, we carried out robustness analyses to determine whether inclusion of high-risk homozygous ε4ε4 carriers affected our results using the same models described above. In these analyses the between-subjects factor was ε4-allele carrier status (none vs. any)..

#### 2.5.3 APOE group differences in spatial navigation performance and its relationship to object-location memory

We tested whether the *APOE* effect previously observed in the full sample of *n*=60 participants persisted in this smaller subset of participants who also completed the memory task (*n*=37). We did so by running general linear models with *APOE*, sex and age on the spatial navigation outcome measures. Dependent variables were mean wayfinding distance and border bias in in the Sea Hero Quest game (Coughlan *et al*., 2019). We also tested for an association between spatial navigation and object-location memory by running Pearson correlations, supplemented with Bayesian analyses.

### 2.6 Data availability

Summary data for precision memory metrics and spatial navigation are available <u>through</u> the Open Science Framework (memory: https://osf.io/42sp9/; spatial navigation: https://osf.io/6adqk/). The code for Bayesian mixture modelling with the MemToolbox can be obtained through http://visionlab.github.io/MemToolbox/ (Suchow *et al*., 2013). Code for mixture modelling using a maximum likelihood estimation implemented by Paul Bays and colleagues is available at https://www.bayslab.com/toolbox/index.php (Bays *et al*., 2011).

## 3. Results

A summary of the memory performance metrics as a function of *APOE* group is shown in Table 2. Fig. 3 shows memory and spatial navigation performance by genotype.

**Table 2.**
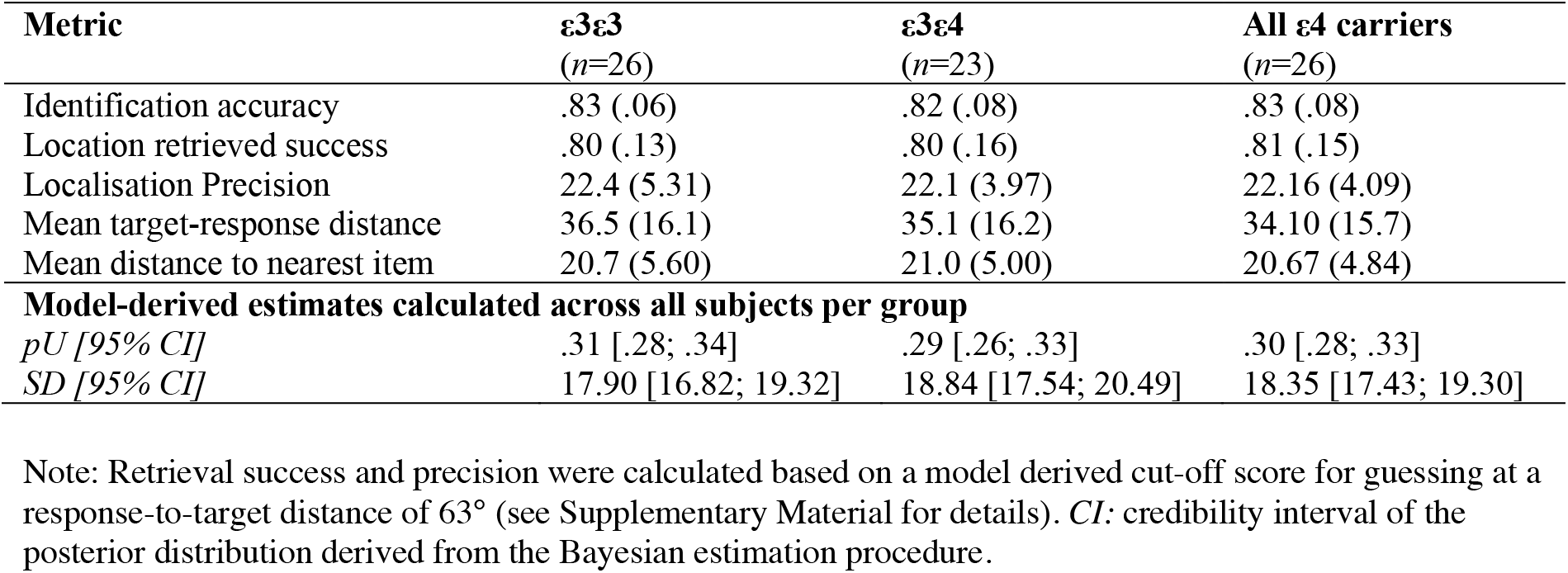
Summary of memory performance across subjects by *APOE* genotype.

**Figure 3.**
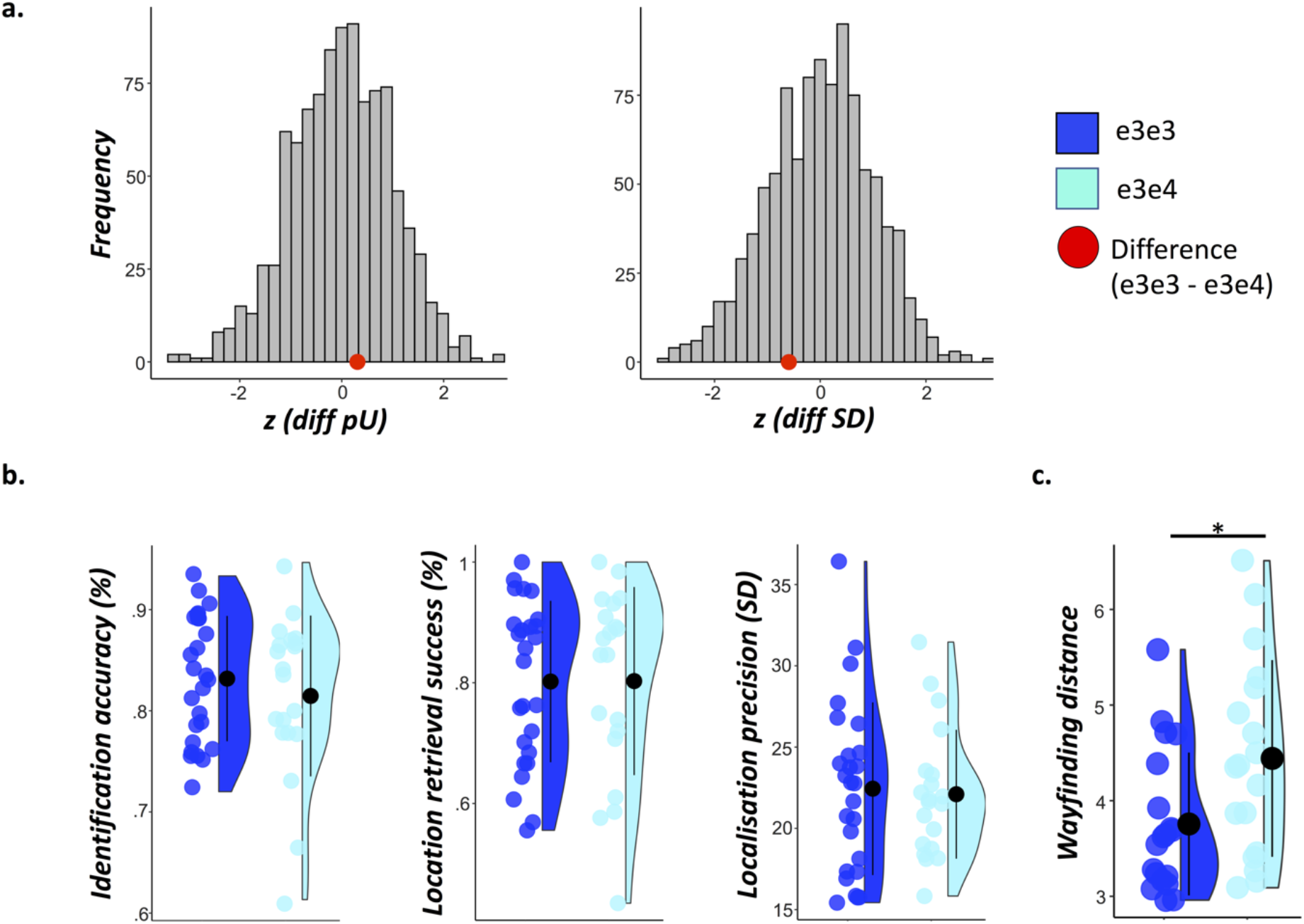
Precision memory and spatial navigation performance by *APOE* genotype. (a) Distribution of standardised group differences derived from 1000 permutations where *n*=26 subjects were randomly assigned to one sample and *n*=20 subjects to another (to match the actual group sizes in our sample). Retrieval success and precision were obtained using mixture modelling on all trials across subjects for the ε3ε44 and the ε3ε4 group, respectively. The red dots represent the standardised true differences in model metrics calculated by subtracting the scores of the ε3ε4 from those of the ε3ε3 group (for *guessing*: *z*=.31; for *SD*: *z*=-.5). (b) Mean ± standard deviation of identification accuracy, retrieval success and precision for each *APOE* group. Retrieval success refers to the proportion of trials falling within 63° of the target object. Precision reflects the standard deviation in response-to-target distance for all trials within 63° of the target object. The *APOE* effect on memory scores and spatial navigation is assessed using general linear models and Bayesian analysis. (c) Mean ± standard deviation of wayfinding distance in the Sea Hero Quest game (Coughlan et al., 2019). \**p*<.05

### 3.1 Group differences based on memory metrics derived from modelling across subjects by APOE group

The results of the permutation analysis are shown in Fig. 3A. The error distributions across subjects in each *APOE* group exhibited considerable overlap. The distribution of permutation-based group differences derived from random assignments to groups confirmed that guessing and imprecision was equivalent in the two *APOE* groups (*guessing*: *z*=.31, *p*=.704=; *imprecision*: *z*=-.59, *p*=.555).

### 3.2 Group differences based on single-subject memory metrics

The linear models controlling for age and sex found no significant effect of *APOE* on identification of objects (*F*_(1,42)_=1.14, *p*=.292, *f*^*2*^=.03, BF_01_=2.17), retrieval success for object locations (*F*_(1,42)_=.45, *p*=.506,, *f*^*2*^=.01, BF_01_=3.02), the precision of recreating locations of correctly retrieved items (*F*_(1,42)_=.12, *p*=.726, *f*^*2*^<.01, BF_01_=3.19), or the mean absolute angular disparity between targets and responses across all trials (*F*_(1,42)_=0.12, *p*=.729, *f*^*2*^<.01, BF_01_=3.37).

### 3.3 Misbinding errors and study-test delay

E4-carriers did not commit more misbinding errors (*F*_(1,42)_=.83, *p*=.367, *f*^*2*^=.02, BF_01_=2.54) or exhibited accelerated forgetting as a function of study-test delay (*F*_(1,42)_=.02, *p*=.890, *f*^*2*^<.01, BF_01_=3.37). All null results held even after inclusion of ε4 homozygotes (Supplementary Material).

### 3.4 Effects of APOE ε4 on spatial navigation

In line with the findings from the full sample in Coughlan and colleagues (2019), among participants who completed both the memory precision and the Sea Hero Quest task ε3ε4 carriers had a longer mean wayfinding distance than ε3ε3 carriers (*F*_(1,33)_=5.60, *p*=.024, *f*^*2*^=.17, *BF*_*10*_=3.44). ε3ε4 carriers in our sub-sample also showed a significant the border bias, although the Bayes Factor did not quite reach the required cut-off of 3 (*F*_(1,33)_=4.54, *p*=.041, *f*^*2*^=.14, *BF*_*10*_=2.55), as it did in the original larger sample (*BF*_*10*_=4.22).

Neither wayfinding distance, nor border bias significantly correlated with retrieval success, precision, mean absolute localisation error, or swap errors (all *r*<.25, *p*>.1). However, the Bayesian analysis could not establish clear support for the null hypothesis for the absence of associations between the object-location memory and spatial navigation performance metrics (all 0<BF_10_ <3).

## 4. Discussion

In this study, we tested whether the precision of long-term memory for object-location binding is affected in healthy middle- and older-aged *APOE* ε4-carriers who do not exhibit impairments on standard neuropsychological tests. We used a continuous report paradigm in which participants were asked to recreate object locations as precisely as possible (Richter *et al*., 2016) and employed Bayesian mixture modelling to separate memory retrieval success from the precision of retrieved locations (Bays *et al*., 2011; Suchow *et al*., 2013). We hypothesised that the precision task combined with mixture modelling may be capable of identifying subtle changes in memory fidelity in preclinical *APOE* ε4-carriers. Previously, preclinical ε4 homozygotes at high risk of AD were impaired on a similar long-term memory fidelity task, while heterozygotes were not (Zokaei *et al*., 2019). Here, we aimed to increase sensitivity of such continuous report paradigms by increasing the to-be-recalled information per test display and by separating memory precision from retrieval success. We then tested if these adjustments may be capable of picking up subtle differences between controls and a genetic risk group, even if the risk group was comprised of individuals with moderate genetic risk of AD (ε4 heterozygotes), around half of whom are expected to develop the disease (Corder *et al*., 1993).

However, we found robust evidence for the absence of an effect of the ε4-allele on object-location long-term memory performance in middle-aged and older adults, regardless of whether the risk groups included only ε4 heterozygotes or additionally added the ε4 high-risk homozygotes. Carriers of the ε4-allele did not recall fewer locations of objects, nor was the precision of their retrieved object-location associations affected. Ε4-carriers also did not commit more misbinding errors of item identity and location. There was no evidence for accelerated forgetting in ε4-carriers as opposed to non-carriers. To our knowledge, this is the first study comparing cognitively healthy *APOE* genotype groups, while using a mixture modelling approach to separate retrieval success from retrieval precision in a task with study-test delays that prevented the involvement of short-term memory. Intriguingly, despite this absence of spatial memory deficits, the ε4-carriers in this sample did exhibit altered wayfinding trajectories in real-time while navigating a virtual environment in the Sea Hero Quest game (Coughlan *et al*., 2019). Moreover, performance on object-location memory and spatial navigation were unrelated.

Few studies have previously investigated memory fidelity in individuals at higher risk for AD during preclinical stages of the disease. Preclinical individuals with higher AD risk based on biomarkers or the *APOE* genotype have been reported to show poorer performance in mnemonic discrimination (combined group of heterozygotes and homozygotes) and continuous report tasks of feature binding in long-term memory (homozygotes) (Liang *et al*., 2016; Sinha *et al*., 2018; Berron *et al*., 2019). Specifically, they exhibit a greater tendency to falsely label as old novel lures that are similar to studied stimuli (Sheppard *et al*., 2016; Sinha *et al*., 2018; Berron *et al*., 2019). They also have higher rates of misbinding, larger object localisation errors (Zokaei *et al*., 2019) and exhibit accelerated forgetting (Zokaei *et al*., 2019; Pavisic *et al*., 2020).

These prior findings suggest that both, aspects of mnemonic discrimination and precision of relational binding may be sensitive to early AD. However, comparisons between these two tasks in terms of their relative sensitivity to AD risk cannot be made at this point given the differences in samples of studies with these tasks in terms of age, neuropsychological deficits, proportion of ε4 heterozygotes and homozygotes and presence of AD pathology (Liang *et al*., 2016; Sheppard *et al*., 2016; Sinha *et al*., 2018; Berron *et al*., 2019, Leal *et al*., 2019*a*; Maass *et al*., 2019; Zokaei *et al*., 2019). Based on one prior study, performance on these two tasks is related and may involve similar but also somewhat dissociable mechanisms (Clark *et al*., 2017). Future studies should aim to compare the sensitivity of mnemonic discrimination tasks and relational binding tasks for the early detection of AD in the same sample.

Based on prior findings of memory fidelity metrics as potentially sensitive markers of preclinical AD, it may be surprising that we did not find an *APOE* effect on memory. However, previous studies have included individuals at higher genetic risk due to presence of the ε4ε4 genotype or familial AD markers (Liang *et al*., 2016; Zokaei *et al*., 2019) or included samples that were on average 5 years older than ours (mean ages of 70 vs. 65 years) and which included neuropsychologically impaired individuals (Sheppard *et al*., 2016; Sinha *et al*., 2018). Our findings therefore suggest that the deficit in object-location memory previously identified could not be detected in individuals that were younger and in a lower genetic risk category, even when using high-sensitivity metrics derived from mixture modelling. As a result, our results do not stand in contrast to prior findings but rather provide information on the potential diagnostic reach of these tasks.

An alternative strategy to test the sensitivity of early detection tasks is to classify cognitively normal preclinical older adults based on tau and amyloid AD biomarker status. To date, this has been done to test for the sensitivity of mnemonic discrimination tests, which show a correlation between both tau and amyloid beta loads with mnemonic discrimination performance (Marks *et al*., 2017; Berron *et al*., 2019, Leal *et al*., 2019*b*; Maass *et al*., 2019; Webb *et al*., 2020). Mean ages in these samples (70+) tend to be significantly older than the participants in the present study (∼65), although in one study the association between tau levels and object mnemonic discrimination could still be observed in individuals aged below 70 years (Berron *et al*., 2019). Interestingly, the association of AD biomarker concentration and mnemonic discrimination deficits remained even after accounting for *APOE* status (Webb *et al*., 2020). Findings from these studies suggest that risk classification based on biomarkers, as opposed to ε4-genotype, may be a better strategy to test the sensitivity of memory precision for early detection of AD in preclinical samples aged 70 or younger without cognitive signs on standard tests (Sperling *et al*., 2020).

Despite the absence of a precision memory deficit in the present sample, Coughlan and colleagues (2019) described suboptimal navigation patterns in these same ε3ε4 carriers 18-months prior to testing (a subset of whom were enrolled in this study). Here, we could reproduce the same wayfinding deficit in the subsample of participants who also completed the precision memory task. This subtle navigational deficit was attributed to a bias toward navigating close to environmental boundaries, as previously documented in an independent cohort (Kunz *et al*., 2015). This very specific impairment may be a result of early tau pathology in the ERC thought to alter the integrity of grid cell representations, which are essential for updating self-motion during navigation (Lithfous *et al*., 2013; Kunz *et al*., 2015; Coughlan *et al*., 2019; Bierbrauer *et al*., 2020; Levine *et al*., 2020). This interpretation is in line with recent evidence suggesting that preclinical ε4-carriers only exhibit spatial navigation deficits in the absence of nearby landmarks or environmental boundaries that normally correct for accumulating temporal error in the grid-cell code (Hardcastle *et al*., 2015; Bierbrauer *et al*., 2020).

Although grid cells are most famously involved in spatial navigation, they also support visual memory (Killian *et al*., 2012). Research suggests that both visual and navigational processes are supported by the entorhinal cortex via common mechanisms that include the formation of spatial or visual maps via grid cells (Nau *et al*., 2018; Bicanski and Burgess, 2019). Specifically, grid cells code for spatial locations in a visual scene much in the same way in which they code for space during exploration of a 3D environment (Killian *et al*., 2012; Nau *et al*., 2018). Based on these findings it has been proposed that grid cells support both spatial navigation and relational memory (Bicanski and Burgess, 2019). It may therefore be surprising that we did not find any effect in our spatial memory precision task and that object-location memory was unrelated to spatial navigation deficits. However, the border bias is a very specific behaviour in ε4-carriers that appears when arenas have larger open spaces where anchoring spatial maps to nearby landmarks cannot be used as a corrective strategy (Kunz *et al*., 2015; Coughlan *et al*., 2019; Bierbrauer *et al*., 2020). Therefore, it has no direct equivalent in 2D visual scene memory in our precision task. This may explain why there is an effect of the ε4-allele on virtual reality spatial navigation but not in object-location memory precision in our sample. A preference for environmental borders may indeed be the very first sign of AD risk dependent behavioural changes, whereas impairment in relational memory may arise at a later stage (Berteau-Pavy *et al*., 2007; Sheppard *et al*., 2016; Sinha *et al*., 2018).

Despite our relatively small sample size, our power analysis suggested that our study had moderate power to detect an *APOE* effect on precision memory similar in magnitude to that that in Coughlan et al. (2019) (Supplementary Material). Even though power was moderate, we could replicate the navigation deficit in this smaller subsample and our Bayesian analysis provided good evidence in favour of a null effect for memory, suggesting that the absence of a genotype effect was not simply due to an inadequate sample size. If a genotype effect on object-location precision does indeed exist in this sample, it is likely to be rather small and may be less meaningful for early detection efforts. This small effect may in part be due to the high heterogeneity of ε3ε4 carriers, given that only a subgroup will move on to actually develop AD (Raber *et al*., 2004). However, the fact that spatial navigations deficits can still be detected even with a small sample as seen here and elsewhere(Kunz *et al*., 2015; Coughlan *et al*., 2019; Bierbrauer *et al*., 2020), suggests that it is indeed possible to find genotype effects on cognition with a sensitive task, even though only 47% of ε3ε4 carriers will move on to develop AD. Our key conclusion, namely that there is no clear object-location memory deficit in ε3ε4 carriers at this age and therefore tests of relational memory may only detect ε4 -dependent deficits at a later point along the AD continuum can still be supported.

To test whether this is indeed the case, it would be informative to follow up the present sample longitudinally to compare participants who do or do not subsequently exhibit cognitive decline associated with AD. Additionally, as discussed above, a promising strategy to test the sensitivity of the precision task in preclinical cases in future studies may be to use biomarkers for classifying individuals into risk groups. This would not only allow studies to determine the sensitivity of memory fidelity metrics but to also assess the specificity of these tasks to AD-related cognitive decline. This is particularly important given the high heterogeneity of ε4-carriers and MCI patients. To date there is still a lack of studies on memory fidelity that stratify MCI patient groups based on AD biomarkers (Troyer *et al*., 2012; Koppara *et al*., 2015; Mowrey *et al*., 2016; Lu *et al*., 2020).

Finally, we argue that it is unlikely that the null findings for object-location memory can be explained on the basis of antagonistic pleiotropy where middle-aged ε4-carriers still have an advantage over ε3ε3 carriers or could stave off the presence of early AD pathology. This explanation is supported for short-term object-location memory (Zokaei *et al*., 2017, 2020; Lu *et al*., 2020). However, it may be less applicable in the case of our results in a task that is more reliant on long-term memory processes and the medial temporal lobe (Berteau-Pavy *et al*., 2007; De Blasi *et al*., 2009; Haley *et al*., 2010; Wolk and Dickerson, 2010; Greenwood *et al*., 2014; Emrani *et al*., 2020). Large-scale studies and meta-analyses across the lifespan have called into question the antagonistic pleiotropy hypothesis in the case of long-term memory (Salvato, 2015; O’Donoghue *et al*., 2018; Weissberger *et al*., 2018; Henson *et al*., 2020) (Salvato, 2015; Weissberger *et al*., 2018; Henson *et al*., 2020). There is only little support for an ε4-dependent advantage in young age (Stening *et al*., 2016) but none for midlife (G. *et al*., 2016), and by older age (comparable to the age in our sample), homozygotes exhibit greater localisation errors than ε3ε3 carriers (Zokaei *et al*., 2019). These prior studies suggest that any potential positive effects of the ε4-allele on spatial memory tasks similar to our object-location paradigm in young adulthood may not carry into late midlife. The effects of the ε4-genotype on long- and short-term memory may unfold differently across the lifespan and we deliberately designed our task to tap into long-term retention processes for which the prodromal hypothesis of *APOE*-ε4 is a more likely explanation.

To our knowledge this is the first study to employ a modelling approach to separate episodic memory retrieval success and precision and test the sensitivity of mnemonic fidelity metrics to preclinical AD risk as measured in a contrast of ε3ε3 and ε3ε4 carriers. Prior work in high risk AD individuals (familial, ε3ε4/ε4ε4, tau and amyloid positive cases) has suggested that object-location memory fidelity may be a sensitive marker for preclinical AD cases and that this effect can be detected in samples aged 70 and older (Liang *et al*., 2016; Sheppard *et al*., 2016; Sinha *et al*., 2018; Zokaei *et al*., 2019; Webb *et al*., 2020). We provide robust evidence that this may not be the case for middle-aged ε3ε4 carriers who were, on average, five years younger than individuals in prior studies. The sensitivity of memory fidelity tasks may therefore not extend to ε4 heterozygotes in their early to mid-60s. Despite no *APOE* genotype effect on object-location memory precision, ε3ε4 carriers in the same sample did exhibit subtle behavioural deficits in spatial navigation. These results provide further support to prior proposals that spatial navigation may be a sensitive marker for the earliest AD-dependent cognitive changes, even before episodic memory (Kunz *et al*., 2015; Coughlan *et al*., 2018). More research in preclinical AD is needed to confirm this hypothesis by direct comparisons of memory fidelity and spatial navigation tasks.

## Acknowledgements

The authors would like to thank their funders and all volunteers who have participated in this study.

## Data availability statement

Data are available at https://osf.io/42sp9/

## Abbreviations

ACE: Addenbrookes Cognitive Examination
AD: Alzheimer’s disease
*APOE*: apolipoprotein
BF: Bayes Factor
ERC: entorhinal cortex
MCI: mild cognitive impairment
pU: proportion of incorrectly remembered trials as estimated in the mixture model approach
pT: proportion of correctly remembered trials as estimated in the mixture model approach
ROCF: Rey-Osterrieth Complex Figure
SD: standard deviation

