## Supplementary Table for "Memory precision of object-location binding is unimpaired in *APOE* ε4-carriers with spatial navigation deficits"

**Supplementary Information**

### 1. Precision memory task

#### *1.1 Task development*

We designed the precision memory tasks according to the following rationale: 1) we aimed to include a number of trials that would allow for a reliable estimate of memory precision and retrieval success, while 2) ensuring that the duration of the task was not too long as to disqualify the task as a potential usable test in a clinical context. A priori, we had set the time limit for a useful potential screening tool at 30 minutes and therefore reduced the number of trials. We piloted multiple versions of the task weighing duration against the total number of trials and arrived at the final task version with 75 trials after running simulations to establish the required task performance that would allow for reliable model estimates of retrieval success and precision, and confirming after a piloting phase that participants' performance did indeed fit the criteria required for reliable model estimation. In the simulations, we tested a range of potential model parameter combinations (guess rates between 5% and 95%; kappa between 2 and 40) and compared the observed parameters that served as model inputs and the expected model parameters that the model fitting procedure would estimate after sampling from the distributions specified by the observed model parameters. Where expected and observed parameter estimates diverged, we could conclude that a task design with the specified number of trials could not reliably estimate mixture model components if mean task performance across participants fell beyond the estimates that could be recovered. The simulations showed that guess rates of up to 55-60% could be tolerated before expected and observed kappa parameters diverge significantly. This is due to the fact that with increasing guess rate, fewer trials are available for the estimation of the von Mises distribution that corresponds to memory precision. The simulations also showed that kappa values between 4 and 40 allowed for reliable estimation of the guess rate as long as guessing does not exceed 60-70% of trials. The pilot data confirmed that the 5 x 5 task version fulfilled these criteria as participants' performance metrics typically lay within these bounds (<60% guessing, kappa between 4 and 40).

#### *1.2 Materials*

Stimuli for the precision memory task consisted of a total of 75 pairs of everyday objects obtained from the Konklab image repository (https://konklab.fas.harvard.edu) and through Google Image Search (Mountain View, CA) and 25 background images (also Google). Each object pair consisted of two different exemplars of the same kind of object (e.g. racket, book). For each pair, one was randomly determined to be the 'target', while the other was chosen to be a 'foil' item. Only the target objects were used to create displays for the object-location memory task study phase. Backgrounds were chosen to display uniform patterns. Three target objects were randomly combined with one background image to create a stimulus display. Objects were placed on pseudo-random locations on an invisible circle centred in the middle of the background image. The randomisation procedure was constrained to ensure no bias in the positions of objects to avoid systematically influencing responses. Separation between objects was a minimum of 62.04 degrees to ensure that objects would not overlap. Displays were identical for all participants but the order of presentation at study and test, respectively, was randomised across subjects.
The experiment was run on Psychtoolbox-3 (http://psychtoolbox.org) on MATLAB.

***1.3** Procedure*

The memory task began with a practice phase where participants saw three displays with three objects each during study and were asked to identify and locate the total of nine objects during the test phase. The experimenter emphasised the importance of paying attention to the features of objects as well as their location. They also stressed that participants should aim to recreate the original position of the test objects as precisely as possible during the location task. In case volunteers felt uncertain about the task instructions after the first training phase, they could choose to complete more practice trials.

Participants received feedback during training and only moved on to the main task after a minimum of five practice displays and after the experimenter had verified good comprehension of the task. After successful completion of the practice phase participants moved on to the study phase for the main task. The task consisted of five blocks each including a study and test phase. During study, participants viewed five displays in a row for eight seconds each. Encoding displays were separated by a fixation cross, which appeared for one second. The study phase was followed by an interference task where participants were asked to count backwards in multiples of three from a random number between 50 and 100 for 12 seconds in order to prevent rehearsal of memory content before the test phase. Participants then completed 15 test trials. For each of the encoding displays, all three objects were tested in sequence. A test trial began with the *identification* question where the target object was presented next to the corresponding foil item on a white background. Participants pressed the *'1'* key to endorse the item on the left-hand side of the screen as old and the *'2'* key to choose the right-hand item. If participants chose the correct item, they moved on to the *location* question where the chosen object appeared on the corresponding encoding display in a random location on a white dial centred around the midpoint of the display. In the middle of the dial the word "Location" printed in white cued participants to the objective of the task. Participants used the arrow keys to move the object clockwise (arrow pointing to the right) or counter-clockwise (arrow to the left) around the dial. There was no time limit, but participants were encouraged to respond within 15 seconds before the location cue turned red in order to keep response times relatively comparable between participants. Participants logged their response by pressing the space bar.

### 2. Mixture modelling of error responses

Note that our primary interest was in comparing the ε3ε3 and ε3ε4 groups as to avoid the admixture of different genotype groups. However, we also provide the reader with information regarding the contrast of ε3ε3 carriers and all carriers of the ε4-allele.

*2.1 Assumptions of models fit to localisation errors*

We fit three models to the data (see Fig. 2A):

1) Model 1 assumes that participants always remember the (approximate) locations of all presented items and may simply vary with respect to the trial-by-trial distance between target and responses. In this case, their responses could best be modelled by using a circular Gaussian (von Mises) distribution, the full-width half-maximum of which represents the precision of a subject's responses.

2) Model 2 assumes that participants may recall the location of the target item for some trials with varying degrees of error but in other trials may forget the location of the target object entirely and resort to guessing [1,2]. In this case, responses can best be characterised in terms of a uniform distribution reflecting random guesses and by a von Mises distribution reflecting the precision of remembered trials. Prior studies in both young and older adults have shown that this model best captures memory responses at longer delays in tasks comparable to the current paradigm [3,4].
3) Model 3 assumes that in addition to random guesses and correct recalls, participants also mistake the location of a non-target item in the same studied display for that of the target item. In this case they would place the probe object where another object had previously been presented, committing a binding error. Misbinding can be modelled by von Mises distributions centred at the location of one of the two non-target items.

Input to the models were the angular disparity between response and target locations for Models 1-3 and additionally for Model 3 the angular disparity between the target and the two non-targets, respectively, to account for potential misbinding errors.

*2.2 Choice of method for model estimation and model selection*

Most prior studies on memory precision have used one of two methods for mixture modelling: the MemToolbox (MT) developed by Suchow and colleagues [5] or the mixture model code written by Paul Bays (https://www.paulbays.com/code.php). The MemToolbox provides the options of parameter estimation with either a Bayesian framework or Maximum Likelihood Estimation, whereas the Bays code employs the latter approach

The mixture modelling approach using the Bays code has previously been used by studies in our lab [3,4]. Other recent work on the effects of the *APOE* genotype on memory has used the MemToolbox for model estimation [6]. We decided to maximise comparability with this study from Zokaei and colleagues on the effects of the *APOE* genotype on short-term location memory precision by using the Bayesian approach from the MemToolbox [5,6]. The Bayesian approach determines a probability distribution for the model parameters (such as for the uniform and von Mises distributions, respectively), showing which parameters could reasonably describe the response data. The posterior distribution is found using a Markov Chain Monte Carlo algorithm. The Maximum A Posteriori estimates (the peak of the posterior distribution) for guessing and *SD* were taken as measures of retrieval success and precision for the models fit to all ε3ε3, all ε3ε4 participants, and all ε4-carriers, respectively. Plots of the posterior distribution by APOE genotype are shown in Supplementary Fig. 1.

We used Akaike, Bayesian and Deviance Information Criteria (*AIC*, *BIC*, *DIC*) for model selection. Supplementary Table 1 shows a summary of the model comparison metrics. The best model was the Standard Mixture Model containing both a uniform distribution for guessing and a von Mises distribution to represent target-response disparity within correctly retrieved trials.

**Supplementary Table 1. Model fit metrics.**

| Model comparison | *AIC* | *BIC* | *DIC* | Best model *AIC* | Best model *BIC* | Best model *DIC* |
|---|---|---|---|---|---|---|
| *ε3ε4* | | | | | | |
| Model 2 – Model 1 | -552.38 | -545.15 | -552.51 | Model 2 | Model 2 | Model 2 |
| Model 2 – Model 3 | -2.00 | -9.23 | -1.56 | | | |
| *ε4ε4* | | | | | | |
| Model 2 – Model 1 | -342.90 | -335.96 | -343.06 | Model 2 | Model 2 | Model 2 |
| Model 2 – Model 3 | -2.00 | -8.95 | -1.19 | | | |
| *All ε4-carriers* | | | | | | |
| Model 2 – Model 1 | -897.78 | -889.99 | -897.33 | Model 2 | Model 2 | Model 2 |
| Model 2 – Model 3 | -2.00 | -9.79 | -0.87 | | | |

**Notes**. Model 1: Von Mises = Correct responses; Model 2: Von Mises + uniform = Correct responses and guessing; Model 3: Von Mises + uniform + von Mises for non-targets = Correct responses, guessing and misbinding errors. In the model comparisons fit metrics are always subtracted from Model 2 such that negative values favour Model 2.

The Bays and MT methods use different parameters to denote the frequency of correct recall and the precision of correctly recalled information. MT uses the proportion of guess responses (*g*), which is equivalent to *pU* as in the work by Bays and colleagues. This can be reverse coded such that higher values reflect better performance by calculating retrieval success as *pT*=1-*pU*. The MT provides the parameter *SD* as a measure of imprecision, reflecting the standard deviation of the von Mises distribution fit to responses coded as degrees between the target and the response. This has the advantage of being easily interpreted because it reflects variability in the location response in native circular space. In contrast, the Bays code uses the concentration parameter kappa (*K*) of the von Mises distribution to describe precision, which is derived from fitting the model to responses expressed in radians. Higher values reflect higher precision.

To allow for comparison with prior work from our lab, we also converted the standard deviation metric that expressed location precision as the concentration parameter of the von Mises distribution which is denoted by *K.* We show both model metrics (*K* and *SD*) in the Supplementary Table 2 below [2,5].

**Supplementary Table 2. Model metrics for the Standard Mixture Model (uniform + von Mises) by APOE status.**

| Group | *pT* | *K* | *g* | *SD* |
|---|---|---|---|---|
| *ε3ε3 carriers* | .69 | 10.77 | .31 | 17.90 |
| *ε3ε4 carriers* | .71 | 9.78 | .29 | 18.83 |
| *All ε4 carriers* | .70 | 10.28 | .30 | 18.35 |

**Note**. *K* and *pT* were derived from code by Paul Bays using maximum likelihood estimation, whereas *SD* and *g* (identical to *pU* used throughout the main article) were obtained using the MemToolbox Bayesian modelling method.

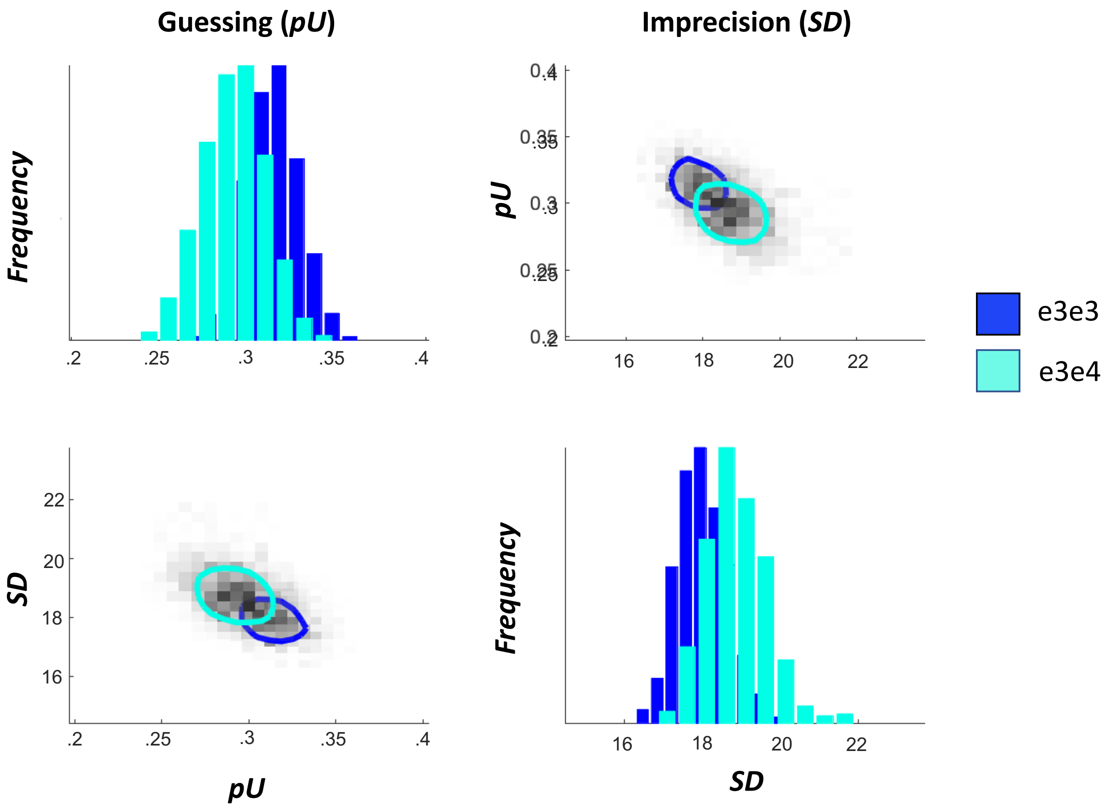


**Supplementary Figure 1. Direct comparison of posterior distributions of model parameters for ε3ε3 and ε3ε4 groups.** The posterior distribution was obtained during the fitting procedure for Model 2 and its maximum (the *maximum a posteriori* estimate) was used as the model parameter estimate for precision and retrieval success, respectively.

#### *2.3 Convergence of Markov Chains*

We used the inbuilt MemToolbox convergence function to visually assess whether the chains in the Markov Chain Monte Carlo (MCMC) approach for the standard mixture model converged. For each *APOE* group, the chains converged well to provide an estimate of precision and the proportion of guessing (Supplementary Fig. 2).

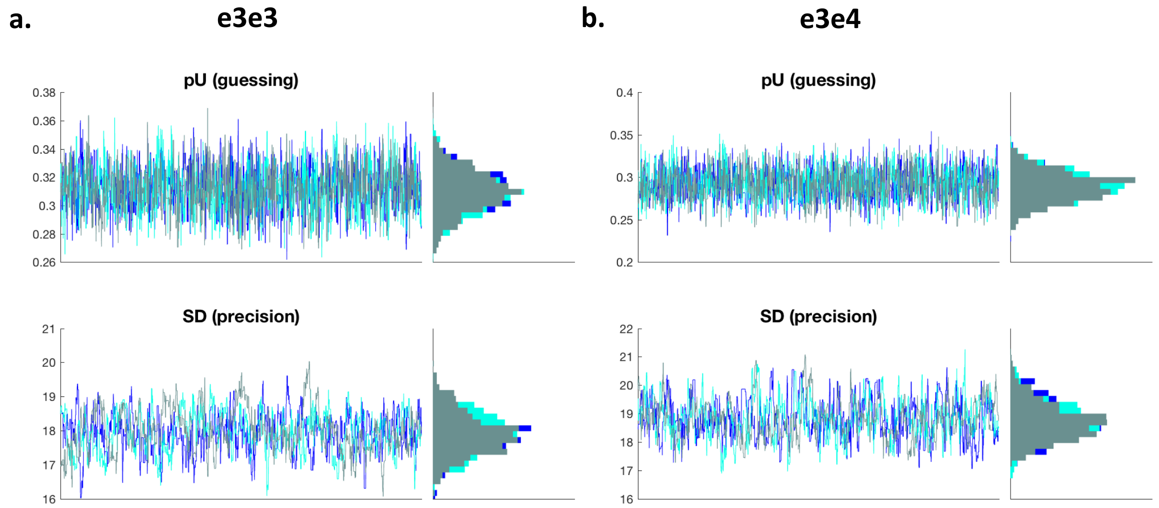


**Supplementary Figure 2. Plot of MCMC chain convergence by APOE genotype group.**

We then also tested whether single-subject responses could be modelled to obtain retrieval success and precision metrics. As we expected, the chains of the model fitting procedure converged well for the majority of participants in our sample, but in four cases, the procedure failed completely. This typically occurs in individuals who exhibited low retrieval success. This can be explained based on the small number of correctly retrieved trials that can be used to estimate the precision parameter. A comparison of good and poor model estimation can be seen in Supplementary Figure 3A and 3B, respectively.

A

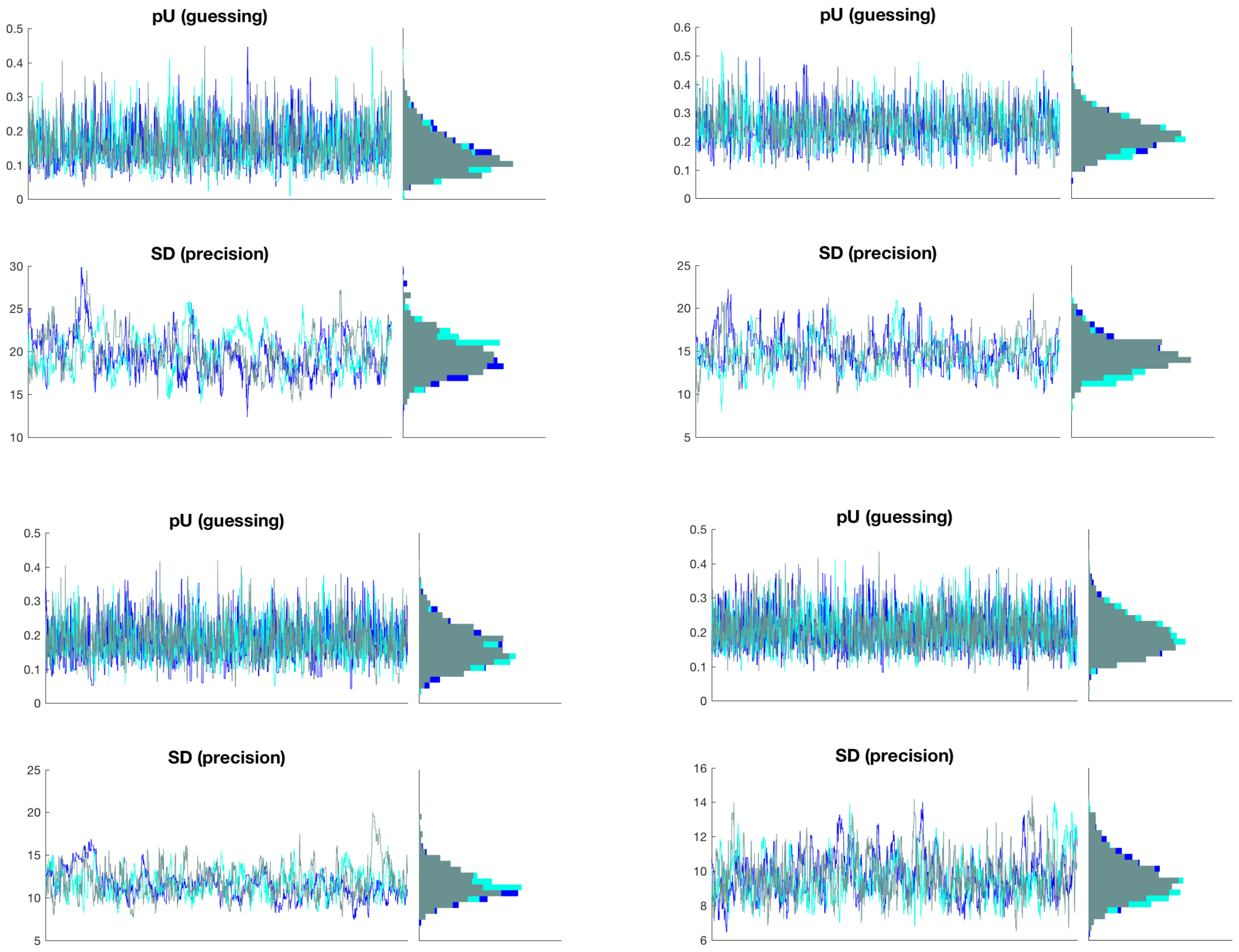
pU (guessing)
SD (precision)
pU (guessing)
SD (precision)
pU (guessing)
SD (precision)
pU (guessing)
SD (precision)

B

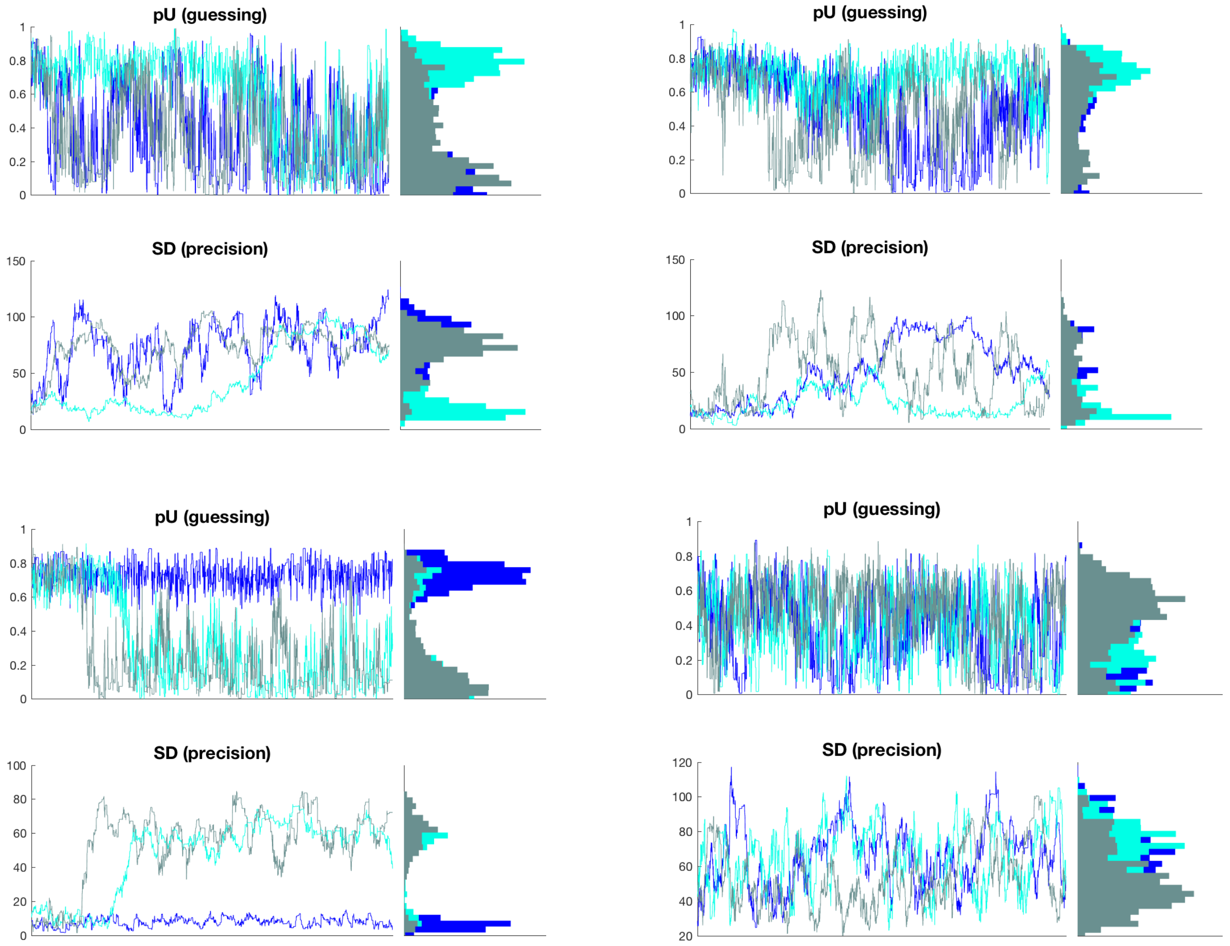
pU (guessing)
0
0.2
0.4
0.6
0.8
1
SD (precision)
0
50
100
150
pU (guessing)
0
0.2
0.4
0.6
0.8
1
SD (precision)
0
50
100
150
pU (guessing)
0
0.2
0.4
0.6
0.8
1
SD (precision)
0
20
40
60
80
100
pU (guessing)
0
0.2
0.4
0.6
0.8
1
SD (precision)
20
40
60
80
100
120

#### *3. Calculating alternative estimates of precision and retrieval success for each subject*

As discussed above, for the vast majority of participants, the chains converged well and model estimates had reasonable confidence intervals. These data confirmed our prior findings from the pilot data showing that model fitting worked well in the majority of cases but may fail for some participants with poor performance. Our task design therefore lends itself to adequate model estimation in most participants.

Given our sample size we decided to avoid data loss to retain as much power as possible. We therefore did not exclude participants with poor model fit as. This decision is justified given that these participants were not extreme outliers on the mean localisation error metric meaning that they could still be included in analyses that do not rely on single subject model estimation.

To allow for estimates of retrieval and precision for individual participants, we followed a procedure previously used by mixture modelling studies to identify trials within the von Mises distribution [4,7,8]. Instead of defining an individual degree cut-off for guessing for every single subject as was done in the mixture modelling approach, we determined a guessing cut-off based on the combined sample of ε3ε3 and ε3ε4 subjects by fitting the standard model to responses across trials. We were then able to determine the distance between target and response at which trials were highly likely to reflect a random guess.

Because the two genetic groups had uneven sample sizes which may have biased the model estimates towards metrics from the larger group (i.e. ε3ε3 carriers), we calculated the cut-off using a permutation procedure over 1000 iterations where a subset with $n$=20 subjects from the ε3ε3 group was chosen to match the $n$=20 subjects from the ε3ε4 group. In each permutation, all trials from the total of 40 subjects in the two groups were used to calculate the probability that a given response in a given trial was a guess (uniform distribution) or a correct response (von Mises distribution). This was done by running the *CO_16* function from Paul Bays' mixture modelling code on the localisation data across all trials from the 40 randomly assigned subjects (https://www.paulbays.com/code.php). This function provides a probability estimate for each trial to determine the likelihood that the response belonged to the uniform or the von Mises distribution. These probabilities were used to calculate the degree value that represents the boundary between successfully and unsuccessfully remembered objects.

For each of the 1000 permutation samples, we determined which response-to-target distance corresponded to a 5% chance of belonging to the von Mises distribution and used this distance as cut-off to identify guessed trials. We then took the average of the guessing cut-offs across all permutations (63°) to classify each trial as forgotten or remembered. For each subject, we computed retrieval success as the proportion of correctly remembered trials as those trials where the target-to-response distance was equal to or smaller than the derived cut-off of 63°. The standard deviation of the distances between response and target across all correctly remembered trials was used to measure of precision, where larger values reflected poorer precision.

#### *4. Obtaining study-test delay data*

Prior work has demonstrated an interaction between study-test delay and the APOE ε4-allele on short-term memory versions of continuous object-location tests with ε4-carriers at an advantage at short delays of 1s which subsides at longer delays beyond 4s (Zokaei *et al.*, 2017, 2020). Here we tested whether this ε4-dependent steeper performance decline as a function of delay extends to our long-term retention task. Displays were not shown in the same order in the study and the corresponding test phase. We therefore calculated the delay between the first presentation of an object during the study phase and its appearance in the location question in the test phase. In this manner, a study-test delay could be obtained for each tested object. For each participant we correlated the study-test delay with the absolute localisation error. Next, we used regressions and Bayesian analysis to test whether performance decreased more steeply with longer time delays in APOE e4-carriers compared to ε3ε3 carriers.

***5. Inclusion of*** ε4 ***homozygotes.***

Even with the addition of the high risk ε4 homozygotes the comparison between ε4-carriers and non-carriers showed no effect of AD genetic risk on object identification ($F_{(1, 45)}$=.22, $p$=.641, $f^2$<.01, $BF_{01}$=3.16), retrieval success ($F_{(1, 45)}$=.08, $p$=.774, $f^2$<.01, $BF_{01}$=3.33), precision of object location memory ($F_{(1, 45)}$=.06, $p$=.808, $f^2$<.01, $BF_{01}$=3.35), mean absolute localisation error ($F_{(1, 45)}$<.001, $p$=.999, $f^2$<.01, $BF_{01}$=3.36) or the occurrence of swap errors ($F_{(1, 45)}$=.33, $p$=.569, $f^2$<.01, $BF_{01}$=3.09). The association between study-test delay on localisation errors was not dependent on *ε4*-carrier status ($F_{(1, 45)}$=..04, $p$=.840, $f^2$<.01, $BF_{01}$=3.31). Looking at the performance of each ε4ε4 carrier individually, none of the *Z*-scores of the ε4ε4 carriers computed on the basis of the ε3ε3 group were larger than ±1.80 in any of item identification, retrieval success, precision, misbinding errors or effect of study-test delay, indicating no significant impairment in any of the participants at the highest genetic risk.

***6. Power analysis to determine issues of underpowering.***

The paper by Coughlan et al. (2019) found a large negative effect of the APOE ε4-allele on wayfinding distance in a virtual spatial navigation task in this sample, 18 months prior to the collection of the memory precision data. Based on the *t*-test the authors conducted, the effect size for this difference was large (Cohen's *d*=.82, corresponding to partial eta squared of .14). Our sample size would allow the detection of an effect of that magnitude in the precision memory task with 76% power if tested with a *t*-test without covariates.

In Table 3 we show which memory scores of the ε4-carriers would correspond to large and moderate effect sizes assuming that the scores of the ε3-carrier group and standard deviations of both groups remained stable. We also calculated the minimum effect size that could be detected with 70% power as *d*=.76 and show in the rightmost column of Table 3. The true effect sizes in our sample are remarkably small and as a result the required scores to achieve large or moderate effects of genotype would have to deviate significantly from the scores observed in our sample.

**Supplementary Table 3. Means and standard deviations of true scores and hypothetical scores that correspond to large and moderate effect sizes for the effect of genotype in the object-location precision task.** The rightmost column corresponds to scores for an effect size of *d*=.76, the effect we could detect with 70% power.

| **Metric** | **True scores** **ε3ε3** (*n*=26) | **True scores** **ε3ε4** (*n*=20) | **True *d*** | **Scores *d*~.82 ε3ε4** (*n*=20) | **Scores *d*~.50 ε3ε4** (*n*=20) | **Scores *d*~.76 with 70% power ε3ε4** (*n*=20) |
|---|---|---|---|---|---|---|
| Identification accuracy | .83 (.06) | .82 (.08) | .14 | .77 | .79 | .78 |
| Location retrieval success | .80 (.13) | .80 (.16) | <.01 | .68 | .72 | .69 |
| Localisation Precision | 22.44 (5.31) | 22.10 (3.97) | .07 | 26.36 | 24.93 | 25.99 |
| Mean target-response distance | 36.46 (16.12) | 35.08 (16.20) | .09 | 49.71 | 44.86 | 48.73 |
| Mean distance to nearest item | 20.69 (5.60) | 20.98 (5.00) | .05 | 25.08 | 23.47 | 24.76 |

When using regression analyses with three predictors (as was done in our case using age, sex and APOE genotype), our sample would allow us to detect a meaningful $R^2$ change as a result of adding the final predåictor (APOE status) with 70% power. The analyses were carried out using G*Power 3.1 for Mac [10].

These power calculations are only appropriate for the analyses based on Frequentist statistics. Establishing null effects using Frequentist statistics is problematic and it has been argued that the only conclusion from a null test based on a *p*-value can be that the effect size was not different enough from 0 to confirm the alternative hypothesis without staying below the predetermined error rate of $\alpha = .05$.[11] Indeed, the $f^2$ effect sizes we found for item identification and item location memory as indexed by a change in $R^2$ between the control model and the model with APOE genotype were so small that sample sizes needed to establish significance of these effects with 80% power are so large as to call into question the practical relevance of the object-location memory test vis-à-vis the spatial navigation task used previously by Coughlan and colleagues.[9] Required sample sizes to detect a change in $R^2$ between our tested models are 264, 787, 2619, and 395 for item identification, retrieval success, precision or mean absolute error, and swap errors, respectively.

### *7. References*


1. Zhang W, Luck SJ. Discrete fixed-resolution representations in visual working memory. 2008;453(May):233-236. doi:10.1038/nature06860
2. Bays PM, Catalao RFG, Husain M. The precision of visual working memory is set by allocation of a shared resource. *J Vis*. 2011;9(10). doi:10.1167/9.10.7.The
3. Korkki SM, Richter FR, Jeyarathnarajah P, Simons JS. Healthy Ageing Reduces the Precision of Episodic Memory Retrieval. *Psychol Aging*. 2020;35(1):124-142. doi:10.1037/pag0000432
4. Richter FR, Cooper RA, Bays PM, Simons JS. Distinct neural mechanisms underlie the success , precision , and vividness of episodic memory. Published online 2016:1-18. doi:10.7554/eLife.18260
5. Suchow JW, Brady TF, Fougnie D, Alvarez GA. Modeling visual working memory with the MemToolbox. *J Vis*. 2013;13(10):1-8. doi:10.1167/13.10.9
6. Zokaei N, Grogan J, Fallon SJ, et al. Short-term memory advantage for brief durations in human APOE ε 4 carriers. *Sci Rep*. Published online 2020:1-10. doi:10.1038/s41598-020-66114-6
7. Korkki SM, Richter FR, Simons JS. Hippocampal-cortical encoding activity predicts

the precision of episodic memory. *bioRxiv*. Published online 2020:1-38. doi:10.1101/2020.11.10.376814
8. Cooper RA, Ritchey M. Cortico-hippocampal network connections support the multidimensional quality of episodic memory. *Elife*. 2019;8:1-37. doi:10.7554/elife.45591
9. Coughlan G, Coutrot A, Khondoker M, Minihane AM, Spiers H, Hornberger M. Toward personalized cognitive diagnostics of at-genetic-risk Alzheimer's disease. *Proc Natl Acad Sci*. 2019;116(19):201901600. doi:10.1073/pnas.1901600116
10. Faul F, Erdfelder E, Lang AG, Buchner A. G*Power 3: A flexible statistical power analysis program for the social, behavioral, and biomedical sciences. *Behav Res Methods*. 2007;39(2):175-191. doi:10.3758/BF03193146
11. Harms C, Lakens D. Making 'null effects' informative: statistical techniques and inferential frameworks. *J Clin Transl Res*. Published online 2018:1-24. doi:10.18053/jctres.03.2017s2.007
